# Methylation-Based Age Estimation in a Wild Mouse

**DOI:** 10.1101/2020.07.16.203687

**Authors:** Tom J. Little, Aine N. O’Toole, Andrew Rambaut, Tamir Chandra, Riccardo Marioni, Amy B. Pedersen

## Abstract

The age structure of populations, or the ageing rate of individuals, impacts aspects of ecology, epidemiology and conservation. Yet for many wild organisms, age is an inaccessible trait. In many cases measuring age or ageing rates in the wild requires molecular biomarkers of age. Epigenetic clocks based on DNA methylation have been shown to accurately estimate the age of humans and laboratory mice, but they also show variable ticking rates that are associated with mortality risk above and beyond that predicted by chronological age. Thus, epigenetic clocks are proving to be useful markers of both chronological and biological age, and they are beginning to be applied to wild mammals and birds. We have acquired strong evidence that an accurate clock is possible for the wood mouse *Apodemus sylvaticus* by adapting epigenetic information from the laboratory mouse (*Mus musculus*). *Apodemus sylvaticus is* a well-studied, common small mammal in the UK and Europe, which is amenable to large-scale experimental perturbations and longitudinal sampling of individuals across their lives. These features of the wood mouse system offer opportunities to disentangle causal relationships between ageing rates and environmental stress. Our wood mouse epigenetic clock is PCR-based, and so requires only tiny amounts of tissue accessible through non-destructive sampling. We quantified methylation using Oxford Nanopore sequencing technology and present a new bioinformatics pipeline for data analysis. We thus describe a new and generalizable system that should enable ecologists and other field biologists to go from small tissue samples to an epigenetic clock for their study animal, which will enable investigations of ageing in the wild which where previously inaccessible.

## Introduction

Outside of long-term field studies of typically quite large animals, age is an inaccessible trait for most wild organisms. Yet, age is a powerful predictor of individual condition and performance, enabling predictions of mortality, susceptibility to parasites and autoimmunity conditions, life history patterns, or a population’s capacity to act as a reservoir of infectious agents. Measuring and understanding the age structure of populations is also a key component of animal ecology and conservation,

Some known methods for estimating age require destructive sampling, e.g. the use of eye lenses in mammals (Rowe et al., 1985), limiting their use in ecological studies. Molecular biomarkers of age can overcome this limitation. Accurate biomarkers of age based on DNA methylation (DNAm) have now been developed for humans and laboratory mice, and these hold considerable promise for use in wild populations (De Paoli-Iseppi et al., 2019; Parrott & Bertucci, 2019; Polanowski et al., 2014; Wright et al., 2018). These ‘epigenetic clocks’ can estimate an individual’s age to within a few years for humans and a few weeks for laboratory mice (Chen et al., 2016; Han et al., 2018; Horvath & Raj, 2018). DNAm-based epigenetic clocks also show variable ticking rates that are associated with mortality risk above and beyond that predicted by chronological age (Chen et al., 2016; Marioni et al., 2015). Thus, an epigenetic clock reveals biological age by identifying individuals who are age-accelerated or decelerated compared to the population average. Importantly, epigenetic clocks appear to hit a sweet spot, being a consistent estimator of both age and age acceleration (Jylhävä et al., 2017; Lee et al., 2016; Marioni et al., 2015). For example, epigenetic clocks show an average correlation with human age of R=0.71 (SE = 0.06, average of correlation coefficients from 13 human cohorts (Chen et al., 2016), although two studies with very small chronological age ranges in their study cohort contribute substantially to the variation). Yet recent comparisons of biomarkers of age in humans also concluded that the epigenetic clock outperforms all other approaches in estimating age acceleration, including telomere length (Jylhävä et al., 2017; Lee et al., 2016). Accelerated epigenetic age in humans is related to diet, lifestyle (Levine et al., 2018; Quach et al., 2017), infection (Horvath & Levine, 2015; Kananen et al., 2015) and reproduction (Binder et al., 2018; Levine et al., 2016).

Epigenetic clocks may thus be a powerful and versatile breakthrough that can substantially advance the ecology and evolution of ageing. Here, we provide evidence that an accurate clock will be possible for the short-lived wild wood mouse *Apodemus sylvaticus* by adapting epigenetic information from the laboratory mouse (*Mus musculus*). The wood mouse is a well-studied field system that is amenable to experimental perturbations and longitudinal sampling of individuals across their entire, and relatively short, lifespans. These features are valuable in comparison to studies on humans and other long-lived species, where disentangling the causal relationships between ageing rate and environmental stress is challenging. The development of this wood mouse epigenetic clock made use of a formerly-wild, now laboratory-reared, wood mouse colony (Clerc et al., 2019; Sweeny et al., 2020). Thus, our starting point was a set of wood mice of known ages, but we also discuss how epigenetic clocks can be applied to wild animals where age is unknown. We used a PCR-based, targeted sequencing approach that makes use of tiny amounts of tissue that can be collected by field biologists without destruction of the animal. Lastly, we made use of cutting edge nanopore sequencing technology, and present a detailed bioinformatics pipeline that can easily be adopted by other researchers to enable a wide range of research questions on any wild mammal.

## Methods

We maintain a formerly-wild, but now lab-reared wood mouse colony in standard laboratory conditions at the University of Edinburgh. The colony has been in captivity for many generations, but the wood mice are purposely outbred to maintain genetic diversity. All mice are housed individually in ventilated cages (Techniplast, 1285L) with food and water ad libitum. DNA was extracted using a phenol-chloroform method from ear punches taken from individuals of known age from the colony. We obtained DNA from 48 mice of both sexes, spanning an age range of 88 to 496 days old. This slightly unusual choice of ages arises because we utilised mice that were part of other experiments. Lifespan in wild mice is not known for certain, and many will die from extrinsic mortality (e.g. predation), but in our own field work we recapture 10-20% of tagged mice the following year (Pedersen, unpublished data), meaning wild mice may can live for hundreds of days.

We adapted the methods of Han *et al* (Han et al., 2018) to use polymerase chain reaction (PCR) to identify CpG sites in *Apodemus sylvaticus* that could contribute to a methylation-based epigenetic clock. We first bisulphite treated the DNA using the ZYMO-Gold DNA bisulphite conversion kit. Bisulphite treatment converts unmethylated cytocine to uracil, which are then converted to thymine during PCR. Methylated cytosines are unchanged by the treatment. For PCR of bisulphite-converted DNA, we focused initially on five genes shown by Han *et al* to be good markers of laboratory mouse age (Han et al., 2018): *Prima1, HSP4, Kcns1, Gm9312* and *Gm7325.* From *Ensemble* we obtained *Mus musclus* DNA sequence that included the key sites identified by Han et al (Han et al., 2018). We blasted 200-300bp of *Mus* sequence for each gene into the *Apodemus sylvaticus* whole genome shotgun sequence available on *NCBI* (taxonID: 40375) to retrieve the homologous wood mouse sequence. We then designed *Apodemus-*specific primers for each gene in approximately the same location as that used by Han *et al* (Han et al., 2018). In some cases, *Apodemus* contained a CpG at the desired primer location, but it was usually possible to design primers slightly up or downstream. In two cases, however, we inserted an ambiguous base pair at a CpG location. Primer sequences are listed in *Appendix 1.*

We used an Oxford Nanopore Flongle to sequence our amplicons and determine the % methylation at all CpG sites (Figure 1). The Flongle sequences up to ~ 1GB of DNA, but given the short length of our amplicons this provides considerable depth of coverage. Our molecular methods follow Quick et al 2017 (Quick et al., 2017), with minor modifications. Briefly, amplicons were pooled by mouse and quantified with a Qubit, the aim being to normalise DNA quantities so that ~ 200 fmol of DNA were loaded onto the Flongle. Otherwise, the minion was run according to the manufacturer’s instructions (Ligation Sequencing Kit, SQK-LSK109).

**Figure 1.**
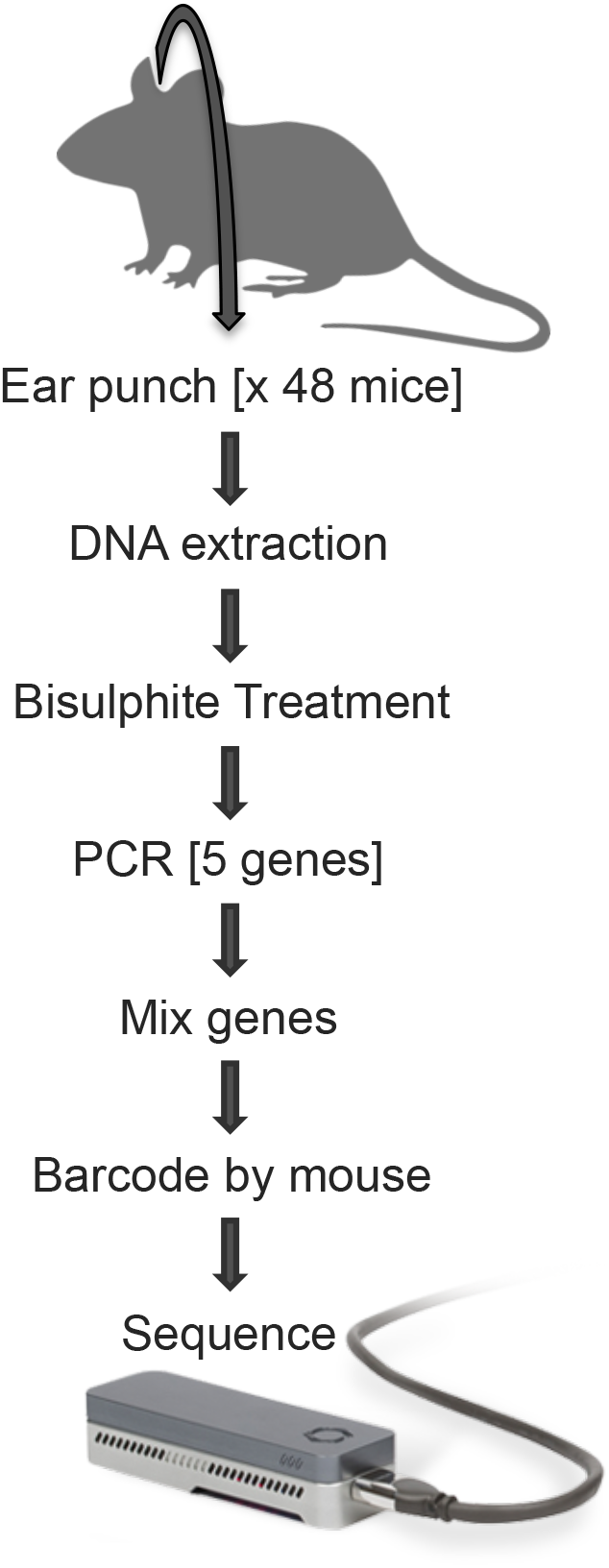
Molecular pipeline from mouse to Flongle for obtaining methylation status at CpGs read from PCR amplification of bisulphite-treated DNA

We developed the custom software ‘Paramether’ on a Snakemake(*20*) framework for easy-to-use estimation of CpG methylation from nanopore sequences. Paramether first bins the sequences by barcode using Porechop (https://github.com/rrwick/Porechop). We curated a database of reference genes representing the bisulphite-treated, amplified gene sequences. Paramether queries this database using parasail (https://github.com/jeffdaily/parasail.git) in order to classify which gene each read corresponds to. Using the alignment with the reference genes, key sites are identified in the read and `Paramether` determines if a site is modified (i.e. a C or a T) or not. This process is continued for each read and counts of methylation are output per gene per barcode.

Proportion methylated Cs at each of the CpGs across the sequences was studied using penalised regression to identify sites that correlate with age. The was achieved with the *GLMNET* package in R (R Core team, 2013) using a LASSO model (mixing parameter alpha=1) and leave-one-out cross validation (nfolds=nrow).

## Results

Four of the five genes (*Prima1, HSP4, Kcns1* and *Gm7325)* studied showed variable methylation at CpG sites, while a fifth (*Gm9312*) showed 100% methylation at all sites and was removed from the analysis. Across these four remaining genes we found 85 CpG sites, and then used a penalised regression to indicate that 9 CpG sites had a probable relationship with age. These sites showed a strong correlation with age (R = 0.85, Figure 2), with a Root Mean Square Error of 59 days.

**Figure 2.**
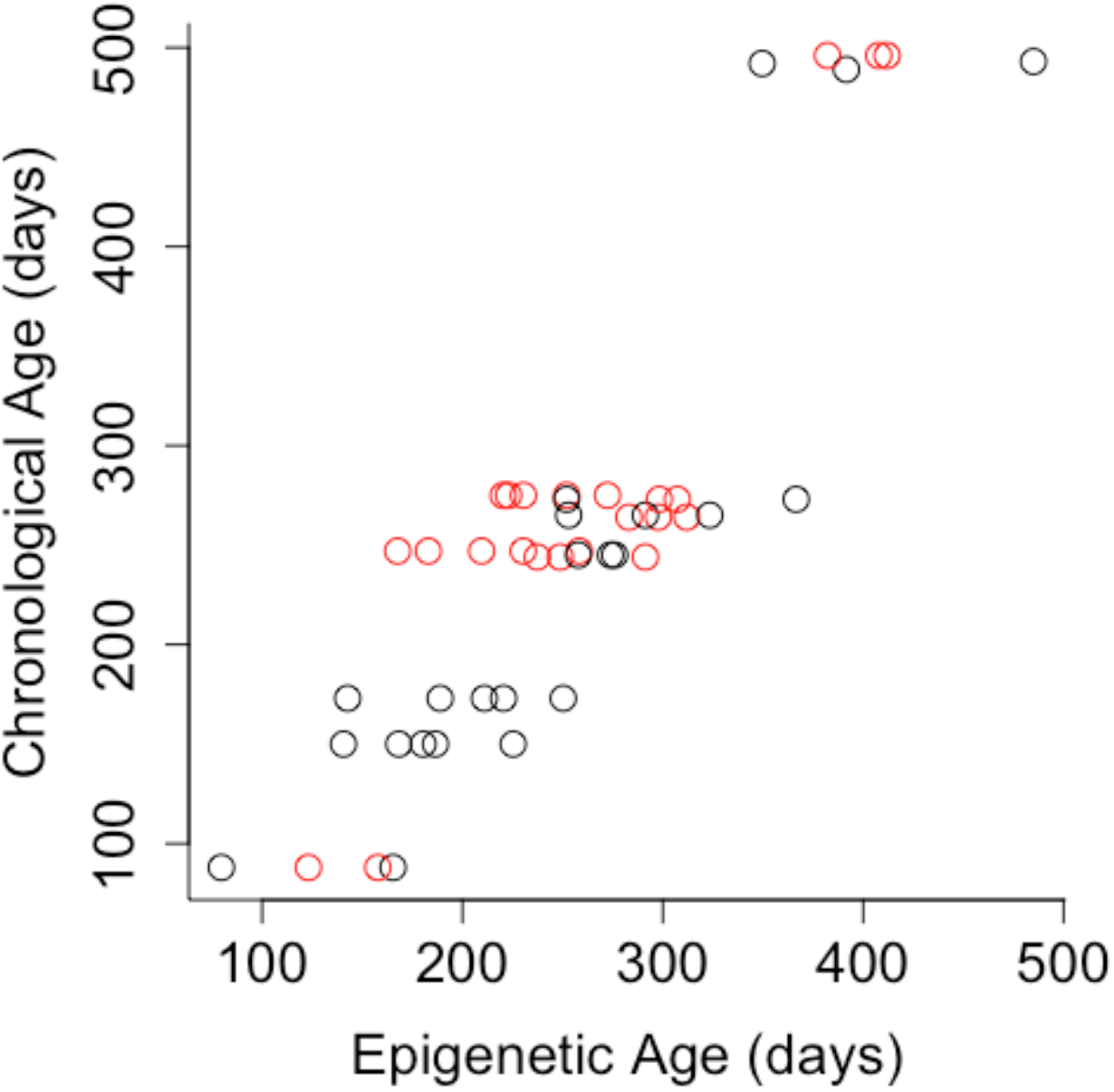
The relationship between age and age predicted by methylation of 4 genes in the wood mouse. R=0.88. The Root Mean Square Error is 58, thus we can identify mouse age to 58 days on average in wood mice in the laboratory. Red=male, black=female

## Discussion

We estimated percent methylation in genes where methylation is associated with age in the laboratory mouse *Mus musclus* (Han et al., 2018). In *Apodemus*, we identified a set of 9 CpGs that show a strong correlation with age (R = 0.85), estimating individual age within a group of mice to ~58 days on average. It would be straightforward to refine our clock using our molecular methods and analysis pipeline to include further genes, for example the remaining genes in Han et al (Han et al., 2018) that we have not yet assessed. While this clock could be improved, the level of accuracy we have achieved using four genes could still prove useful for studies of wild wood mice, whose lifespan ranges from months to a few years at most. More specifically, this clock could determine which mice survived the winter and which mice are newly born in the spring summer, which could have important impcliations for population dynamics. Past work on *Apodemus* has shown that many mice are recaptured within a few months, and some are captured across years. Thus our clock will be applicable given expected recapture rates, enabling the study of epigenetic-based ageing rates in relation to various environmental stressors.

Our results also confirm that methylation may accumulate similarly with age between different rodent species that are >10 million years diverged: *Mus musculus* and *Apodemus sylvaticus* (Steppan et al., 2004). Methylation patterns are not always well conserved across species, especially when they are only distantly related (*22*), but at least within Class, the development of epigenetic clocks appears to benefit from a high degree of conservation (De Paoli-Iseppi et al., 2017; Polanowski et al., 2014; Wright et al., 2018), which could greatly foster the study of new taxa. Still, in many cases it will be necessary, or even desirable (for example to obtain the most informative epigenetic clock possible), to develop clocks through whole-genome approaches (De Paoli-Iseppi et al., 2017, 2019). Regardless of the molecular approach taken to develop epigenetic clocks, the deployment of them to large samples sizes of wild individuals may often require a targeted sequencing approach due to either financial limitations or due to restrictions in the amount or type of tissue that can be collected.

It also seems likely, as we have observed, that clocks developed on laboratory-kept animals will be correlated tightly with age, as individuals living in such stable conditions are unlikely to exhibit much age acceleration or deceleration. Clocks developed directly on wild animals might show weaker associations with age because these samples will contain individuals that have experienced a wider range of ecological conditions and environmental stressors, some of which are likely to impact ageing rate. It is unknown how clocks developed through different approaches will provide information on age versus age acceleration, but we do know that human clocks have proven informative for both. Granted, most human clocks are based on a sizable number of sites (often in the hundreds), which can be analysed with the aid of high throughput methylation arrays. Clocks developed with a comparatively small number of sites have yet to be extensively tested (but see (Olova et al., 2019)), but do require only tiny amounts of tissue, are preparable with existing technology and do not require service facilities (which may not always be available).

A further challenge faced by many study systems, is that deploying a clock to the field means studying individuals of unknown age. Longitudinal sampling of the same individuals will be a key approach ecologists will adopt in testing, for example, the role of environmental factors on age acceleration. To progress this, consider that epigenetic age acceleration is routinely (in human studies) measured as epigenetic age corrected for chronological age. This can be investigated by regressing epigenetic age onto chronological age and then testing whether specific factors, (e.g. nutrition, environmental conditions, or measures of infection or immunity), explain the residual variation, in a statistical model such as: Epigenetic age ~ Age + factor. However, where chronological age is unknown, but individuals can be captured repeatedly, the time lapse between samples, can be substituted for ‘Age’ and the change in epigenetic age becomes the response variable indicating the ageing rate: Δepigenetic age ~ Δage + Factor. Such an analysis can provide a direct estimate of age acceleration in the wild, whilst testing if candidate predictors are drivers of the acceleration.

Thus many of the challenges of applying epigenetic clocks to wild animals seem surmountable, and DNA methylation-based clocks may be superior to any other approach. For example, much past work on ageing in the wild has used telomere lengths as a marker of age (Belsky et al., 2017; Jylhävä et al., 2017). Whilst telomere attrition may differ between long and short-lived species(Foley et al., 2018), within species the relationship between telomere length and age appears weak (Belsky et al., 2017, 2017; Fairlie et al., 2016; Foley et al., 2018; Jylhävä et al., 2017), and their use may be fraught with unresolved technical issues (Belsky et al., 2017; Nettle et al., 2019; Reichert et al., 2017). Telomere lengths may be good indicators of the state of an individual and thus predict mortality, but even this may be less accurate than what can be achieved with epigenetic clocks (Jylhävä et al., 2017). Epigenetic clocks based on DNA methylation are only beginning to be used on wild organisms, but with their potential accuracy for measuring age and their sensitivity to ageing rate, they may well prove to be a boon to the ecology of ageing.

## Data Accessibility

The pipeline output data have been deposited at https://doi.org/10.5061/dryad.vq83bk3qj. Porechop, which we used to bin the sequence reads is available at https://github.com/rrwick/Porechop. Parasail is at https://github.com/jeffdaily/parasail.git

## Notes

### Competing Interest Statement

The authors have declared no competing interest.

